# Development of environmental DNA surveillance for the threatened crucian carp (*Carassius carassius*)

**DOI:** 10.1101/344317

**Authors:** Lynsey R. Harper, Nathan P. Griffiths, Lori Lawson Handley, Carl D. Sayer, Daniel S. Read, Kirsten J. Harper, Rosetta C. Blackman, Jianlong Li, Bernd Hänfling

## Abstract

1. The crucian carp (*Carassius carassius*) is one of few fish species associated with small ponds in the UK. These populations contain genetic diversity not found in Europe and are important to conservation efforts for the species, which has declined across its range. Detection and monitoring of extant crucian carp populations are crucial for conservation success. Environmental DNA (eDNA) analysis could be very useful in this respect as a rapid, cost-efficient monitoring tool.
2. We developed a species-specific quantitative PCR (qPCR) assay for eDNA surveillance of crucian carp to enable non-invasive, large-scale distribution monitoring. We compared fyke netting and eDNA at ponds with (N = 10) and without (N = 10) crucian carp for presence-absence detection and relative abundance estimation, specifically whether DNA copy number reflected catch-per-unit-effort (CPUE) estimate. We examined biotic and abiotic influences on eDNA detection and quantification, and compared qPCR to standard PCR. Notably, eDNA occurrence and detection probabilities in relation to biotic and abiotic factors were estimated using a hierarchical occupancy model.
3. eDNA analysis achieved 90% detection for crucian carp (N = 10), failing in only one pond where presence was known. We observed an overall positive trend between DNA copy number and CPUE estimate, but this was not significant. Macrophyte cover decreased the probability of eDNA occurrence at ponds, whereas CPUE and conductivity had positive and negative influences on eDNA detection probability in qPCR replicates respectively. Conductivity also had a negative effect on DNA copy number, but copy number increased with temperature and percentage of macrophyte cover. PCR was comparable to qPCR for species detection and may provide semi-quantitative information.
4. Our results demonstrate that eDNA could enable detection of crucian carp populations in ponds and benefit ongoing conservation efforts, but imperfect species detection in relation to biotic and abiotic factors and eDNA workflow requires further investigation. Nonetheless, we have established an eDNA framework for crucian carp and sources of imperfect detection which future investigations can build upon.

## 1. Introduction

The crucian carp (*Carassius carassius*) (Figure 1) is a cryptic, benthic fish species popular with anglers (Copp, Warrington & Wesley, 2008b; Sayer et al., 2011). As one of few fish associated with small ponds, this species may have an important ecological role but its relationship with other lentic biodiversity is understudied (Copp & Sayer, 2010; Stefanoudis et al., 2017). Although listed as ‘Least Concern’ on the International Union for Conservation of Nature (IUCN) Red List of Threatened Species, the species has declined throughout its native range of Northwest and Central Europe (Copp et al., 2008b; Sayer et al., 2011), with local extinctions across the UK (Copp & Sayer, 2010). The county of Norfolk in eastern England was believed to hold abundant and widely distributed crucian carp populations, but research indicates heavy (~75%) declines in this region (Sayer et al., 2011). Declines of the crucian carp throughout its range are due to habitat loss (Copp et al., 2008b; Sayer et al., 2011), species displacement by the invasive gibel carp (*Carassius gibelio*) (Copp et al., 2008b; Tarkan et al., 2009; Sayer et al., 2011), and genetic introgression through hybridisation (Hänfling et al., 2005). Indeed, Sayer et al. (2011) observed only 50% of crucian carp ponds were not inhabited by goldfish (*Carassius auratus*), common carp (*Cyprinus carpio*), or their hybrids with crucian carp.

**Figure 1.**
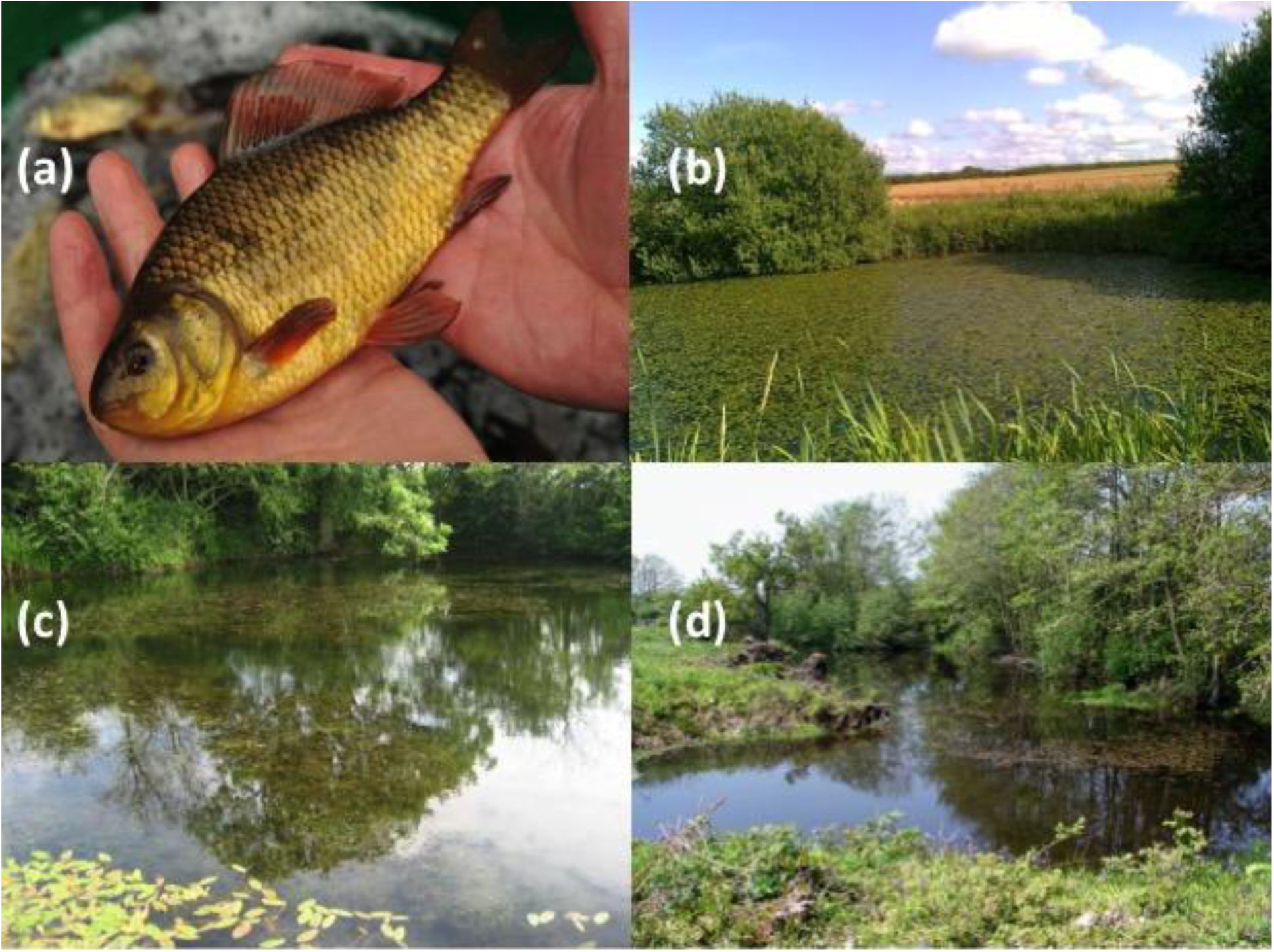
Crucian carp (*Carassius carassius*) individual **(a)** and examples of ponds inhabited by crucian carp **(b-d)**. Photo **(a)** by John Bailey.

Prior to the 1970s, crucian carp were thought to have been introduced to the UK alongside common carp and were classed as non-native (Maitland, 1972). Wheeler (1977) deemed the species native to southeast England based on archaeological evidence and a historic distribution that mirrored native cyprinids. Conservation organisations (e.g. English Nature, Environment Agency) later recognised the crucian carp as native and threatened (Smith & Moss, 1994; Environment Agency, 2003), but recent genetic evidence supports anthropogenic introduction of the crucian carp to the UK during the 15th century (Jeffries et al., 2017). Nonetheless, many introduced species in the UK are now naturalised, and several provide ecological and economical benefits (Manchester & Bullock, 2000). Evidence suggests the crucian carp is characteristic of species-rich ponds (Copp et al., 2008b; Sayer et al., 2011; Stefanoudis et al., 2017), and English populations contain a substantial proportion of the overall genetic diversity for the species across Europe. These populations may buffer species displacement by gibel carp (Jeffries et al., 2017), but are threatened by hybridisation with goldfish and possible displacement (Hänfling et al., 2005; Tarkan et al., 2009) as well as anthropogenic activity (Copp, Černý & Kováč, 2008a).

In 2010, the crucian carp was designated as a Biodiversity Action Plan (BAP) species in Norfolk (Copp & Sayer, 2010; Sayer et al., 2011). To meet the BAP aims, local conservation efforts have included species reintroduction, pond restoration, and eradication of goldfish (Sayer et al., 2011). However, current distribution records are unreliable as individuals are frequently misidentified as the feral brown variety of goldfish due to physical similarity (Copp et al., 2008a; Tarkan et al., 2009; Sayer et al., 2011), and many populations are mixtures of true crucian carp and crucian carp x goldfish hybrids (Hänfling et al., 2005). Consequently, distribution maps have been called into question and further monitoring is needed to ensure long-term success of established and reintroduced crucian carp populations (Copp et al., 2008a; Tarkan et al., 2009).

Primarily, crucian carp are surveyed using fyke netting or electrofishing but these methods can be costly and time-consuming. Environmental DNA (eDNA) analysis offers a potentially rapid and cost-effective approach to fish monitoring (Jerde et al., 2011; Sigsgaard et al., 2015; Wilcox et al., 2016; Hänfling et al., 2016; Hinlo et al., 2017a). Species are identified using intracellular or extracellular DNA deposited in the environment by individuals via secretions, excretions, gametes, blood, or decomposition (Lawson Handley, 2015). eDNA has been applied worldwide to survey for invasive freshwater fish (Jerde et al., 2011; Keskin, 2014; Robson et al., 2016; Hinlo et al., 2017a), and is now used routinely to monitor Asian carp (*Hypophthalmichthys* spp.) invasion in the Great Lakes, USA (Farrington et al., 2015). A quantitative PCR (qPCR) assay targeting crucian carp was also published in the context of early warning invasion monitoring for fish species that may arrive in Canada (Roy et al., 2017), but was only tested on tissue-derived DNA. Of equal importance to invasion monitoring, eDNA analysis has enhanced surveys for threatened and endangered freshwater fish (Sigsgaard et al., 2015; Schmelzle & Kinziger, 2016; Piggott, 2016; Bylemans et al., 2017).

eDNA analysis has been conducted with conventional PCR (PCR) (Ficetola et al., 2008; Jerde et al., 2011), but qPCR and droplet digital PCR (ddPCR) are suggested to perform better, suffer less from inhibition, and enable abundance or biomass estimation (Nathan et al., 2014). However, these estimates can be inconsistent across habitats and target organisms. In flowing water, Hinlo et al. (2017a) found no relationship between DNA copy number and conventional density estimates of common carp, yet Takahara et al. (2012) observed a positive association between common carp biomass and eDNA concentration in ponds. Environmental variables play a substantial role in abundance/biomass estimation by influencing the ecology of eDNA (Barnes et al., 2014). Variables examined have included temperature, pH, salinity, conductivity, anoxia, sediment type, and UV light (Takahara et al., 2012; Barnes et al., 2014; Pilliod et al., 2014; Keskin, 2014; Strickler, Fremier & Goldberg, 2015; Robson et al., 2016; Buxton et al., 2017b; Buxton, Groombridge & Griffiths, 2017a; Weltz et al., 2017; Stoeckle et al., 2017; Goldberg, Strickler & Fremier, 2018). However, these variables are not always measured and only a handful of studies have assessed their effects in ponds (Takahara et al., 2012; Buxton et al., 2017a, b; Goldberg et al., 2018).

In this study, we developed a species-specific qPCR assay for the threatened crucian carp. We evaluated presence-absence detection with eDNA compared to fyke netting, and whether our assay could estimate abundance by comparing catch-per-unit-effort (CPUE) estimates obtained by fyke netting to DNA copy number. We investigated the influence of biotic and abiotic factors on eDNA detection and quantification, and performed a small-scale comparison of qPCR and PCR for species detection. We hypothesised that: (1) eDNA and fyke netting would provide comparable presence-absence records for crucian carp; (2) DNA copy number would positively correlate with CPUE estimate; (3) eDNA detection and quantification would be influenced by crucian carp density, temperature, pH, conductivity, surface dissolved oxygen, macrophyte cover and tree shading; and (4) qPCR would possess greater detection sensitivity than PCR. We provide an eDNA framework for crucian carp monitoring which holds promise for routine survey.

## 2. Methods

### 2.1 Study sites

We studied 10 ponds with confirmed crucian carp presence at different densities and 10 fishless ponds in Norfolk (Figure 2). This region is low-lying (<100 m above sea level) and mainly agricultural. All ponds were selected to be small (<40 m in max. dimension), shallow (<2 m), macrophyte-dominated, and open-canopy. Ponds were largely surrounded by arable fields, excluding one located in woodland. No specific permits were required for sampling but relevant landowner permissions were obtained.

**Figure 2.**
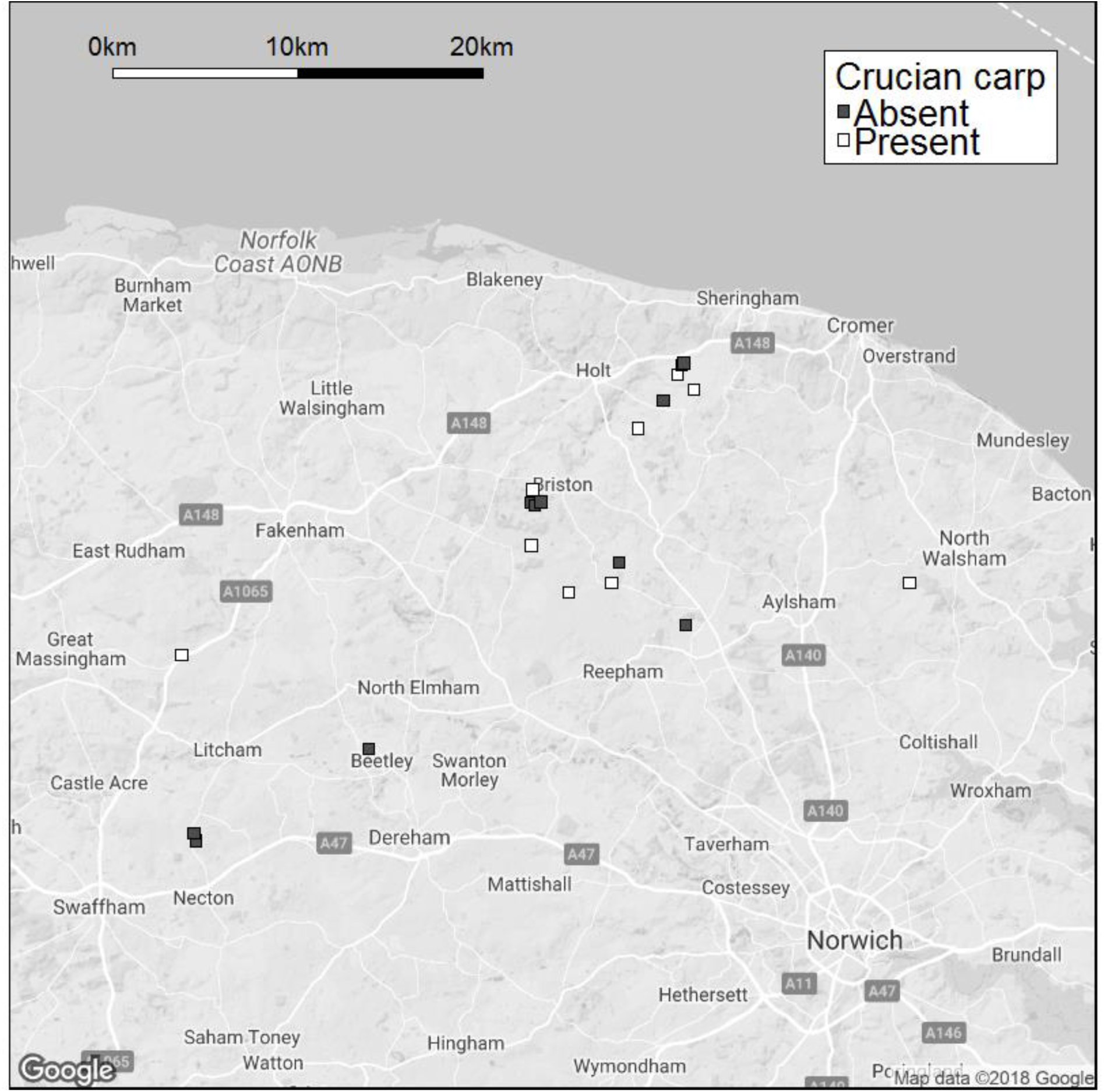
Google map of North Norfolk, eastern England, showing the distribution of ponds stocked with crucian carp (white squares) and ponds where the species is absent (grey squares).

### 2.2. Conventional survey

Crucian carp presence-absence was confirmed at each pond by fyke netting between 2010 and 2016. Bar two ponds surveyed in 2013 and 2015, all crucian carp ponds were last surveyed in 2016. Where possible, double-ended fyke nets were set perpendicular to the bank or to beds of aquatic vegetation and exposed overnight (for c. 16 h), with the number of fyke nets set being proportional to pond size. This provided CPUE estimates of relative densities, which are the number of fish captured per fyke net per 16 h exposure. Environmental data were collected between May and August from 2010 to 2017. Conductivity, pH, surface dissolved oxygen and water temperature were measured with a HACH HQ30d meter (Hach Company, CO, USA), and alkalinity measured by sulphuric-acid titration using a HACH AL-DT kit (Hach Company, CO, USA). Percentage of macrophyte cover and shading (trees and bushes) of ponds were estimated visually.

### 2.3 eDNA sampling, capture and extraction

Five 2 L surface water samples were collected from the shoreline of each pond using sterile Gosselin™ HDPE plastic bottles (Fisher Scientific UK Ltd, UK) and disposable gloves. Samples were taken at equidistant points around the pond perimeter where access permitted. All ponds without crucian carp were sampled on 22^nd^ August 2016. Water samples were transported on ice in sterile coolboxes to the Centre for Ecology and Hydrology (CEH), Wallingford, stored at 4 °C, and vacuum-filtered within 24 hours of collection. Coolboxes were sterilised using 10% v/v chlorine-based commercial bleach solution and 70% v/v ethanol solution before ponds containing crucian carp were sampled on 25^th^ August 2016. Samples were handled in the same way. For each pond, a full process blank (1 L molecular grade water) was taken into the field and stored in coolboxes with samples. Blanks were filtered and extracted alongside samples to identify contamination.

Where possible, the full 2 L of each sample was vacuum-filtered through sterile 0.45 μm cellulose nitrate membrane filters with pads (47 mm diameter; Whatman, GE Healthcare, UK) using Nalgene filtration units. One hour was allowed for each sample to filter but if filters clogged during this time, a second filter was used. After 2 L had been filtered or one hour had passed, filters were removed from pads using sterile tweezers and placed in sterile 47 mm petri dishes (Fisher Scientific UK Ltd, UK), which were sealed with parafilm (Sigma-Aldrich^®^, UK) and stored at −20 °C. The total volume of water filtered and number of filters used per sample were recorded for downstream analysis (Table S1). After each round of filtration (samples and blanks from two ponds), all equipment was sterilised in 10% v/v chlorine-based commercial bleach solution for 10 minutes, immersed in 5% v/v MicroSol detergent (Anachem, UK) and rinsed with purified water.

All filters were transported on ice in a sterile coolbox to the University of Hull and stored at −20 °C until DNA extraction one week later. DNA was isolated from filters using the PowerWater^®^ DNA Isolation Kit (MO BIO Laboratories, CA, USA) following the manufacturer’s protocol in a dedicated eDNA facility at University of Hull, devoted to pre-PCR processes with separate rooms for filtration, DNA extraction and PCR preparation of environmental samples. Duplicate filters from the same sample were co-extracted by placing both filters in a single tube for bead milling. Eluted DNA (100 μL) concentration was quantified on a Qubit™ 3.0 fluorometer using a Qubit™ dsDNA HS Assay Kit (Invitrogen, UK). DNA extracts were stored at −20 °C until further analysis.

### 2.4 Assay design, specificity and sensitivity

We designed a novel assay to target a 118 bp amplicon within the mitochondrial cytochrome *b* (*cytb*) gene, specific to crucian carp. Crucian carp sequences from Jeffries *et al*. (2016) were aligned using MAFFT in AliView (Larsson, 2014) to sequences downloaded from the NCBI nucleotide (nt) database for 24 closely related species of European freshwater fish, and a consensus sequence for each species identified (Figure S1). Sequences were visually compared to maximise nucleotide mismatches between crucian carp and non-target species, particularly goldfish and common carp, and minimise theoretical risk of non-specific amplification. Mismatches in primer regions were maximised over the probe region to increase specificity (Wilcox et al., 2013). Species-specific primers CruCarp_CytB_984F (5′- AGTTGCAGATATGGCTATCTTAA-3′) and CruCarp_CytB_1101R (5′- TGGAAAGAGGACAAGGAATAAT-3′), and corresponding probe CruCarp_CytB_1008Probe (FAM 5′-ATGGATTGGAGGCATACCAGTAGAACACC-3′ BHQ1) were selected on this basis.

Primers without probe were tested *in silico* using ecoPCR (Ficetola et al., 2010) against a custom, phylogenetically curated reference database that was constructed for eDNA metabarcoding of lake fish communities in Windermere, Lake District National Park, England, and contains 67 freshwater fish species confirmed or potentially present in the UK (Hänfling et al., 2016). Parameters set allowed a 50-150 bp fragment and maximum of three mismatches between each primer and each sequence in the reference database. Specificity of primers (without probe) was also tested against the full NCBI nucleotide (nt) database using Primer-BLAST (Ye et al., 2012) with default settings.

Primers were first validated *in vitro* using PCR and tissue DNA (standardised to 1 ng/μL) from fin clips of crucian carp and four closely related non-target species: goldfish, common carp, tench (*Tinca tinca*), and sunbleak (*Leucaspius delineatus*). An annealing temperature gradient (Supporting Information: Appendix 1) was used to ensure optimal amplification of crucian carp and no non-target amplification (Figure S2). The primers were also tested on eDNA samples from ponds recently stocked with crucian carp to confirm potential for eDNA amplification (Figure S3). Molecular grade water (Fisher Scientific UK Ltd, UK) was used as the no template control (NTC) in all tests.

Primer and probe concentrations, standard curve preparation, and cycling conditions for qPCR were then optimised (Supporting Information: Appendix 1). All subsequent qPCR analyses were performed using the conditions detailed in section 2.5. Specificity tests were repeated using qPCR on 10 non-target species related to crucian carp (Table S2, Figure S4) with tissue DNA from fin clips (standardised to 1 ng/μL). The positive control and NTC were crucian carp DNA and molecular grade water (Fisher Scientific UK Ltd, UK) respectively. The limits of detection (LOD, the concentration at which no crucian carp DNA will amplify) and quantification (LOQ, the concentration at which all technical replicates consistently amplify crucian carp DNA) (Agersnap et al., 2017) were established using a 10-fold dilution series of crucian carp DNA (1 to 1 × 10^−8^ ng/μL) and qPCR standards (10^6^ to 1 copy/μL) (Figure S5). Five technical replicates were performed for standards, controls, and samples in tests of assay specificity and sensitivity.

### 2.5 Detection and quantification of crucian carp eDNA

All qPCR reactions were prepared in a UV and bleach sterilised laminar flow hood in our dedicated eDNA facility. Reactions were performed in a total volume of 20 µL, consisting of 2 µL of template DNA, 1 µL of each primer (Forward 900 nM, Reverse 600 nM), 1 µL of probe (125 nM) (Integrated DNA Technologies, Belgium), 10 µL of TaqMan^®^ Environmental Master Mix 2.0 (Life Technologies, CA, USA) and 5 µL molecular grade water (Fisher Scientific UK Ltd, UK). Once eDNA samples and three NTCs were added to each 96-well plate, the plate was sealed and transported to a separate laboratory on a different floor for addition of the standard curve and three positive controls (crucian carp DNA, 0.01 ng/µL) in a UV and bleach sterilised laminar flow hood.

Our standard curve was a synthesised 500 bp gBlocks^®^ Gene Fragment (Integrated DNA Technologies, Belgium) based on GenBank accessions (KT630374 - KT630380) for crucian carp from Norfolk (Jeffries et al., 2016). Copy number for the gBlocks^®^ fragment was estimated by multiplying Avogadro’s number by the number of moles. We performed a 10-fold serial dilution of the gBlocks^®^ fragment to generate a 6-point standard curve that ranged from 10^6^ to 10 copies/µL. eDNA samples were compared to these known concentrations for quantification (Hinlo et al., 2017a). Each standard was replicated five times on each qPCR plate. Similarly, five technical replicates were performed for every sample and full process blank from each pond.

After addition of standards and positive controls, plates were again sealed and transported to a separate laboratory on a different floor where qPCR was conducted on a StepOnePlus™ Real-Time PCR system (Life Technologies, CA, USA). Thermocycling conditions consisted of incubation for 5 min at 50 °C, a 10 min denaturation step at 95 °C, followed by 60 cycles of denaturation at 95 °C for 15 s and annealing at 60 °C for 1 min. We used 60 cycles for consistency with optimisation tests, but cycling could be reduced to 45 cycles for subsequent applications (see Supporting Information: Appendix 1).

Amplifications were considered positive detections if the exponential phase occurred within 45 reaction cycles as the mean C_q_ value was 40.07 for the LOD (1 copy/µL). A pond was considered positive for crucian carp if two or more of the five technical replicates from a sample returned positive, or more than one sample returned any positive technical replicates (Goldberg et al., 2016). False negatives were obtained for one pond therefore all samples were tested for inhibition by spiking duplicate qPCR reactions with a known concentration of crucian carp template (1000 copies/µL) (Jane et al., 2015).

### 2.6 DNA sequencing

Non-target DNA extracts and full-process blanks that amplified with qPCR were Sanger sequenced alongside a representative eDNA sample from each positive pond (N = 9) to confirm sequence identity. Purification and sequencing was performed by Macrogen Europe (Amsterdam, The Netherlands) in triplicate in the forward direction. Sequences were edited using CodonCode Aligner (CodonCode Corporation, MA, USA) with default settings. Sequences were then manually aligned in AliView (Larsson, 2014) and poor quality sequences discarded (Figure S6). Primers were removed from remaining sequences, and sequences identified against the full NCBI nucleotide (nt) database using the NCBI BLASTn tool.

### 2.7 Data analysis

Technical replicates for each qPCR standard that differed by >0.5 C_q_ from the average of the five technical replicates performed were discarded to minimise bias induced by pipetting error. All technical replicates for eDNA samples were retained, and those which failed to amplify were classed as 0 C_q_ (Goldberg et al., 2016). The C_q_ values for each set of technical replicates were averaged and quantified to provide a single DNA copy number for each sample. Samples with no positive amplifications were assigned a DNA copy number of zero. DNA copy numbers of samples were then averaged to generate a single DNA copy number for each pond.

All subsequent data analyses were performed in the statistical programming environment R v.3.4.2 (R Core Team, 2017). Agreement between fyke netting and qPCR for crucian carp detection was assessed using Cohen’s kappa coefficient (Cohen, 1960). Following this, Pearson’s Chi-squared Test for Independence was used to test for difference in the frequency of crucian carp positive and negative ponds between methods. Prior to testing for a relationship between CPUE estimate and average DNA copy number for each pond, we tested normality of the data set using the Shapiro-Wilk test and visually inspected the underlying distribution using histograms. All data points were included as some with the appearance of outliers may be due to environmental fluctuations influencing DNA quality. A Spearman rank correlation coefficient was calculated to measure strength of association as the interval data were not normally distributed. Effects of water volume filtered, number of filters used, and water sample content on DNA copy number of samples were also tested (see Supporting Information: Appendices 1, 2).

The R package ‘eDNAoccupancy’ v0.2.0 (Dorazio & Erickson, 2017) was used to fit a Bayesian, multi-scale occupancy model to estimate crucian carp eDNA occurrence and detection probabilities. Existing eDNA literature was used to identify biotic and abiotic factors reported to affect eDNA detection, persistence and degradation, and construct hypotheses regarding their effects on probability of eDNA occurrence in ponds (ψ), eDNA detection probability in water samples (θ), and eDNA detection probability in qPCR replicates (p). Only macrophyte cover was included as a covariate at site level. Vegetated ponds are more likely to contain crucian carp by offering individuals refuge from predation as well as foraging and spawning opportunities (Sayer et al., 2011), and have reduced UV exposure thereby preserving eDNA (Barnes et al., 2014). Such ponds are susceptible to terrestrialisation which can create anoxic conditions that impede crucian carp reproduction and recruitment (Sayer et al., 2011), although these conditions may slow eDNA degradation and enable detection over longer periods (Barnes et al., 2014; Pilliod et al., 2014; Weltz et al., 2017). At sample level, biotic and abiotic factors were included as covariates. More individuals (reflected by CPUE) should increase eDNA concentration and improve detection in water samples. Temperature can increase physical, metabolic, or behavioural activity of organisms resulting in more eDNA release, breakdown, and degradation (Takahara et al., 2012; Pilliod et al., 2014; Strickler et al., 2015; Robson et al., 2016; Lacoursière-Roussel, Rosabal & Bernatchez, 2016; Buxton et al., 2017b; Bylemans et al., 2017). Links established between eDNA and pH support greater detectability, concentration, and persistence of eDNA in more alkaline waters (Barnes et al., 2014; Strickler et al., 2015; Goldberg et al., 2018). Conductivity relates to Total Dissolved Solids (TDS) and sediment type, which can impair eDNA detection due to release of inhibitory substances and their capacity to bind DNA (Buxton et al., 2017a; Stoeckle et al., 2017). At qPCR replicate level, covariates again included CPUE as higher eDNA concentration should improve amplification success and consistency, whereas conductivity may indicate inhibitory substances that cause amplification failure.

Prior to modeling, all environmental variables were assessed for collinearity using Spearman’s correlation coefficient and Variance Inflation Factors (VIFs) calculated using the R package ‘car’ v2.1-6 (Fox & Weisberg, 2011). Variables were considered collinear and removed if r >0.3 and VIF >3 (Zuur et al., 2009), following which candidate variables were centred and scaled to have a mean of 0. We constructed 64 models which included macrophyte cover at site level, and different covariate combinations at sample and qPCR replicate levels. Models were ranked (Table S3) according to posterior predictive loss criterion (PPLC) under squared-error loss and the widely applicable information criterion (WAIC). The model with the best support was selected for comparison to models without covariates at site and the entire sampling hierarchy.

We examined the influence of abiotic factors on eDNA quantification using a generalised linear mixed effects model (GLMM) within the R package ‘glmmTMB’ v0.2.0 (Brooks et al., 2017). Collinearity was assessed as above, leaving temperature, pH, conductivity, and percentage of tree shading as explanatory variables. Pond was modelled as a random effect to account for spatial autocorrelation in our data set and the influence of other properties inherent to each pond, whereas all other explanatory variables were fixed effects. A Poisson distribution was specified as the nature of the response variable (DNA copy number) was integer count data. Validation checks were performed to ensure all model assumptions were met and absence of overdispersion (Zuur et al., 2009). Model fit was assessed visually and with the Hosmer and Lemeshow Goodness of Fit Test (Hosmer & Lemeshow, 2000) using the R package ‘ResourceSelection’ v0.3-0 (Lele et al., 2014). Model predictions were obtained using the predict() function and upper and lower 95% CIs were calculated from the standard error of the predictions. All values were bound in a new data frame and model results plotted for evaluation using the R package ‘ggplot2’ v2.2.1 (Wickham, 2009).

## 3. Results

### 3.1 Assay specificity and sensitivity

Only crucian carp amplified in ecoPCR, confirming primer specificity. Non-target species returned by primer-BLAST against the full NCBI nucleotide (nt) database were *Barilius bakeri* (a Cyprinid fish restricted to India, 6 mismatches), *Naumovozyma dairensis* (fungi, 8 mismatches), and *Medicago trunculata* (plant, 8 mismatches). Our probe sequence could not be included *in silico* but would likely increase specificity. All crucian carp DNA amplified by PCR, with non-target amplification removed above 58 °C. Tissue extracts from common rudd (*Scardinius erythrophthalmus*) and European chub (*Squalius cephalus*) included in qPCR assay specificity tests were amplified by primers and probe, but possessed low DNA copy number (<10 copies/µL). In a later test, common carp DNA also amplified (<10 copies/µL). However, no amplification was observed for NTCs, fresh tissue extracts obtained from rudd and chub, or eDNA samples from locations where crucian carp were absent and these species were present (data not shown). DNA sequencing confirmed cross-contamination of reference material, where sequences were either identified as crucian carp or poor quality (Table S4). Our assay was highly sensitive with a LOD of 1 copy/µL and LOQ of 10 copies/µL. The lowest concentration of crucian carp tissue DNA that amplified was 0.0001 ng/µL.

### 3.2 qPCR analysis

The qPCR assay had average amplification efficiency of 93.614% (range 79.607-102.489%) and average R^2^ value of 0.998 (range 0.995-0.999) for the standard curve. No amplification occurred in NTCs but the POFA4 full process blank amplified (<10 copies/µL). This was the only contaminated blank as the POHI blank filtered alongside POFA4 and POHI samples, and blanks downstream of these samples did not amplify. Partial inhibition occurred in a single sample from four different ponds: PYES2 (no crucian carp), RAIL, POHI, and GUES1 (crucian carp present). However, amplification of other samples enabled ponds to meet established detection criteria, thus problematic samples were not treated for inhibition or qPCRs repeated.

### 3.3 Presence-absence detection

eDNA surveillance detected crucian carp in 90% of ponds (N = 10) with confirmed presence. Sanger sequencing of representative samples confirmed species identity as crucian carp (Table S5). eDNA failed entirely in one pond (CHIP) that contained a sizeable crucian carp population (CPUE = 60.50), but samples from CHIP were not inhibited. Crucian carp DNA was not detected at any sites where the species was absent. Cohen’s kappa coefficient (κ = 0.9) indicated strong agreement between fyke netting and eDNA analysis, further supported by no significant difference in frequency of crucian carp positive and negative ponds by each monitoring tool (χ^2^ = 0.1003, df = 1, *P* = 0.752).

### 3.4 Relative abundance estimation

We identified a weak, positive trend between CPUE estimate and DNA copy number (Figure 3), but this was not significant (*r_S_* = 0.334, df = 8, *P* = 0.345). The association was unchanged when ponds not surveyed by fyke netting in 2016 were removed, or DNA copy number was set as 10 copies/µL for amplifications below the LOQ.

**Figure 3.**
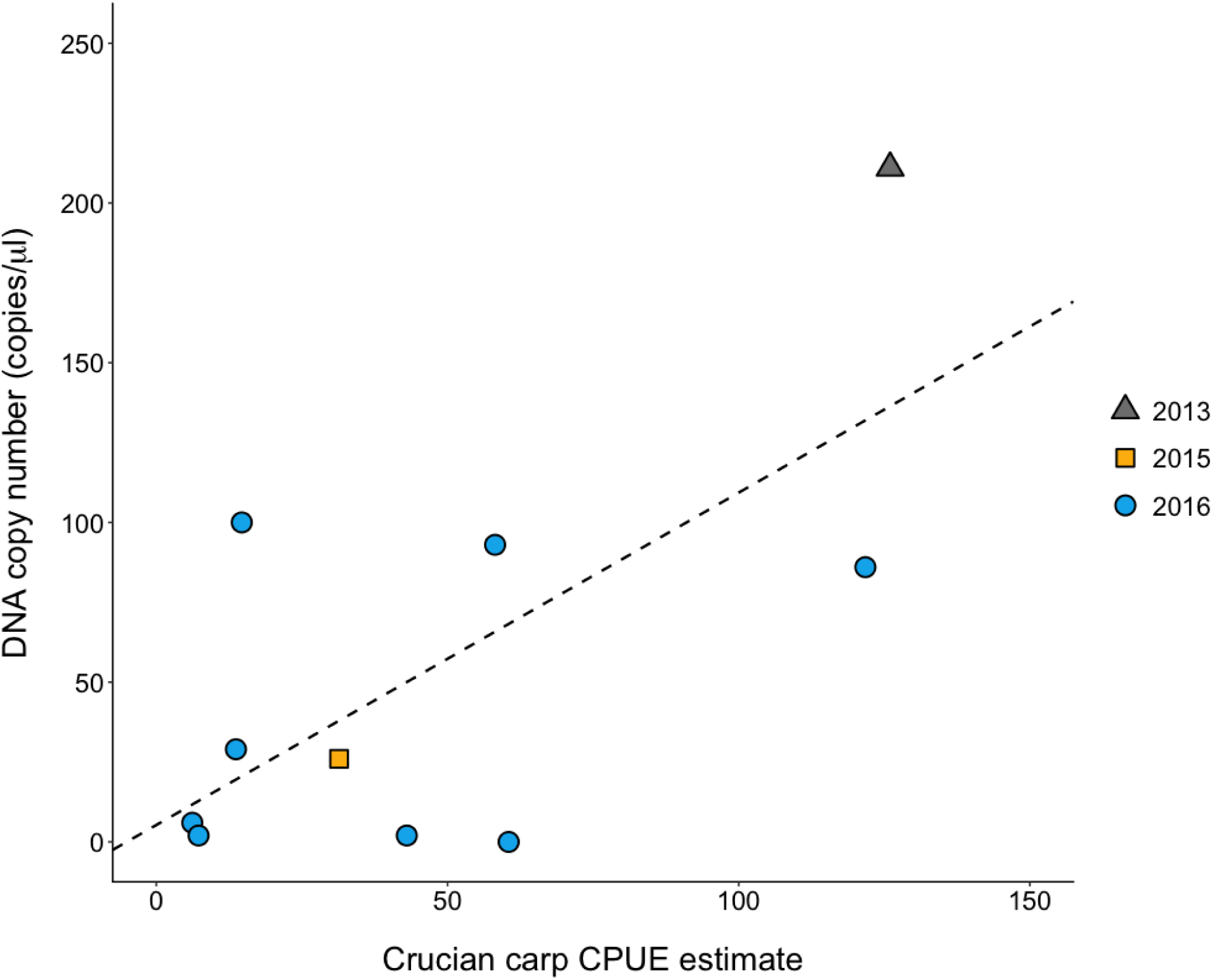
Relationship between DNA copy number and CPUE estimate for crucian carp. A broad positive trend was observed but the relationship was variable, where some ponds with high CPUE had low DNA copy number and vice versa. Points are coloured and shaped by the last year that ponds were surveyed using fyke netting. Three ponds fell below the LOQ (10 copies/μL) and one pond did not amplify during qPCR.

### 3.5 Factors influencing eDNA detection and quantification

The occupancy model with the best support included macrophyte cover as a covariate of eDNA occurrence probability at sites (ψ), and CPUE and conductivity as covariates of eDNA detection probability in qPCR replicates (p). The model did not include any covariates of eDNA detection probability in water samples (θ). The probability of eDNA occurrence in a pond (Figure 4a) ranged between 0.34 to 0.73 (Table 1) and was negatively influenced by macrophyte cover (parameter estimate = -0.294). Estimates of eDNA detection probability in a qPCR replicate ranged between 0.15 to 1.00 (Table 1), where crucian carp CPUE and conductivity played positive (parameter estimate = 1.409) and negative (parameter estimate = -1.923) roles in eDNA availability respectively (Figures 4b, c). The GLMM identified temperature (0.711 ± 0.284, 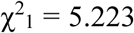, *P* = 0.022), conductivity (−0.006 ± 0.002, 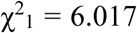, *P* = 0.014), and macrophyte cover (0.035 ± 0.015, 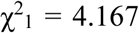, *P* = 0.041) as significant predictors of DNA copy number, where DNA copy number was greater at higher temperatures (Figure 5a) but decreased as conductivity and macrophyte cover increased (Figures. 5b, c).

**Table 1.** Bayesian estimates of crucian carp eDNA occurrence probability at a pond (ψ), eDNA detection probability in a water sample (θ), and eDNA detection probability in a qPCR replicate (p). Posterior median and 95% credible interval (CI) are given for each parameter of the occupancy model. Ponds were all located in Norfolk, eastern England.

| Pond | Crucian carp (Y/N) | CPUE estimate | $\psi$ | | $\theta$ | | $p$ | |
| --- | --- | --- | --- | --- | --- | --- | --- | --- |
|  |  |  | Posterior median | 95% CI | Posterior median | 95% CI | Posterior median | 95% CI |
| CAKE | Y | 43.00 | 0.73 | 0.19 - 0.99 | 0.83 | 0.70 - 0.92 | 0.17 | 0.06 - 0.34 |
| CHIP | Y | 60.50 | 0.43 | 0.23 - 0.65 | 0.83 | 0.70 - 0.92 | 0.16 | 0.05 - 0.40 |
| GUES1 | Y | 121.75 | 0.34 | 0.11 - 0.67 | 0.83 | 0.70 - 0.92 | 0.99 | 0.90 - 1.00 |
| MYST | Y | 6.17 | 0.68 | 0.22 - 0.98 | 0.83 | 0.70 - 0.92 | 0.93 | 0.85 - 0.97 |
| OTOM | Y | 14.67 | 0.37 | 0.15 - 0.65 | 0.83 | 0.70 - 0.92 | 0.96 | 0.90 - 0.99 |
| POFA4 | Y | 13.67 | 0.43 | 0.23 - 0.65 | 0.83 | 0.70 - 0.92 | 0.89 | 0.81 - 0.95 |
| POHI | Y | 7.25 | 0.52 | 0.29 - 0.76 | 0.83 | 0.70 - 0.92 | 0.42 | 0.26 - 0.59 |
| RAIL | Y | 58.17 | 0.37 | 0.15 - 0.65 | 0.83 | 0.70 - 0.92 | 1.00 | 0.99 - 1.00 |
| SKEY1 | Y | 31.38 | 0.37 | 0.15 - 0.65 | 0.83 | 0.70 - 0.92 | 1.00 | 1.00 - 1.00 |
| WADD3 | Y | 126.00 | 0.58 | 0.28 - 0.86 | 0.83 | 0.70 - 0.92 | 1.00 | 1.00 - 1.00 |
| LDUN2 | N | 0 | 0.46 | 0.26 - 0.67 | 0.83 | 0.70 - 0.92 | 0.86 | 0.74 - 0.94 |
| LDUN3 | N | 0 | 0.49 | 0.28 - 0.71 | 0.83 | 0.70 - 0.92 | 0.15 | 0.05 - 0.33 |
| PYES2 | N | 0 | 0.46 | 0.26 - 0.67 | 0.83 | 0.70 - 0.92 | 0.89 | 0.78 - 0.96 |
| SABA | N | 0 | 0.37 | 0.15 - 0.65 | 0.83 | 0.70 - 0.92 | 0.97 | 0.91 - 0.99 |
| VALE | N | 0 | 0.55 | 0.29 - 0.82 | 0.83 | 0.70 - 0.92 | 1.00 | 1.00 - 1.00 |
| WADD10 | N | 0 | 0.40 | 0.19 - 0.65 | 0.83 | 0.70 - 0.92 | 0.31 | 0.15 - 0.50 |
| WADD11 | N | 0 | 0.37 | 0.15 - 0.65 | 0.83 | 0.70 - 0.92 | 0.15 | 0.04 - 0.32 |
| WADD17 | N | 0 | 0.46 | 0.26 - 0.68 | 0.83 | 0.70 - 0.92 | 0.95 | 0.88 - 0.99 |
| WOOD | N | 0 | 0.55 | 0.29 - 0.81 | 0.83 | 0.70 - 0.92 | 0.91 | 0.80 - 0.96 |
| WRONG | N | 0 | 0.34 | 0.11 - 0.67 | 0.83 | 0.70 - 0.92 | 0.93 | 0.83 - 0.98 |

**Figure 4.**
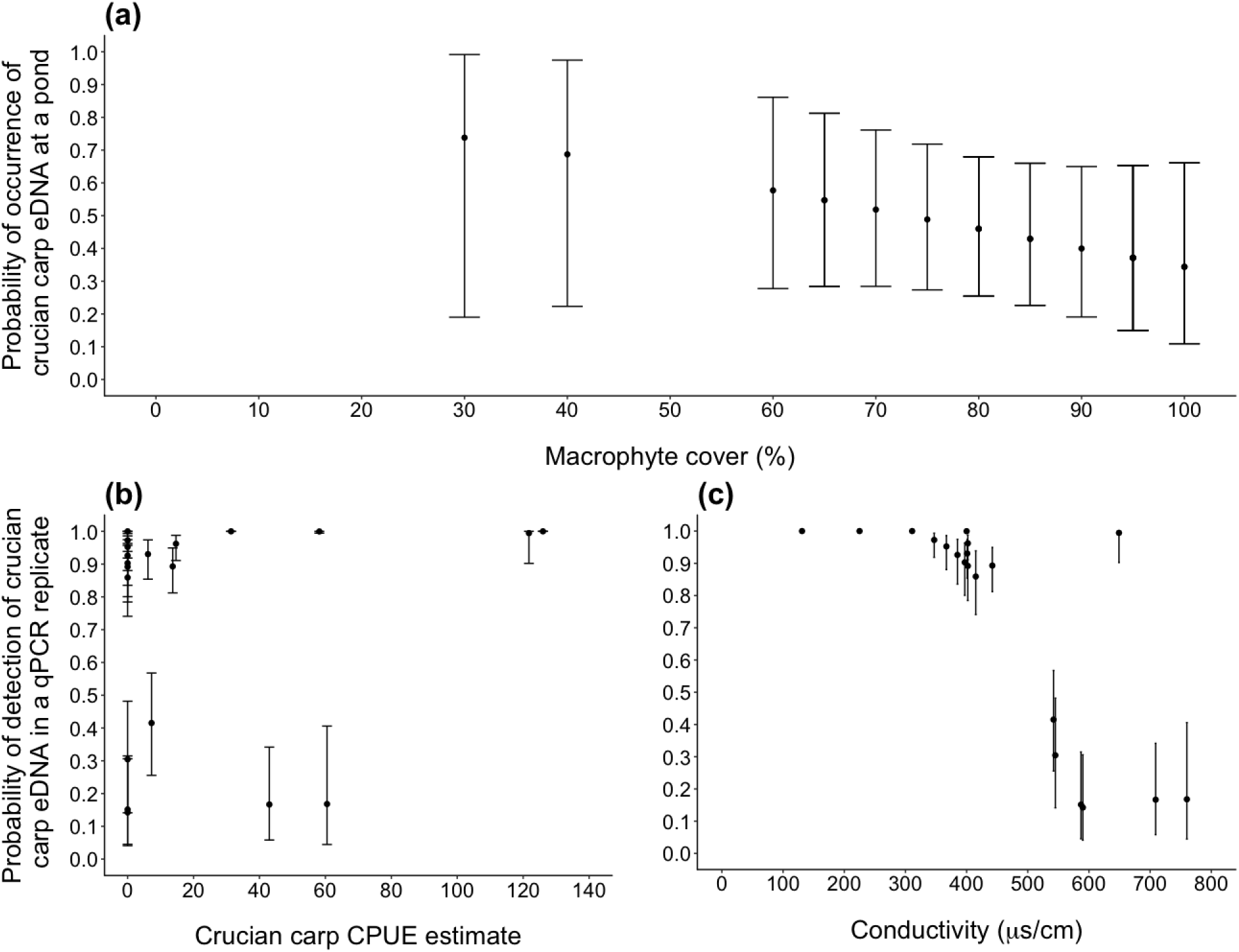
Estimated probabilities of eDNA occurrence in ponds, and eDNA detection in qPCR replicates. Points are estimates of posterior medians with 95% credible intervals. Probability of eDNA occurrence in ponds decreased as percentage of macrophyte cover increased **(a)**. Probability of eDNA detection in qPCR replicates increased with higher CPUE **(b)** but decreased as conductivity increased **(c)**.

**Figure 5.**
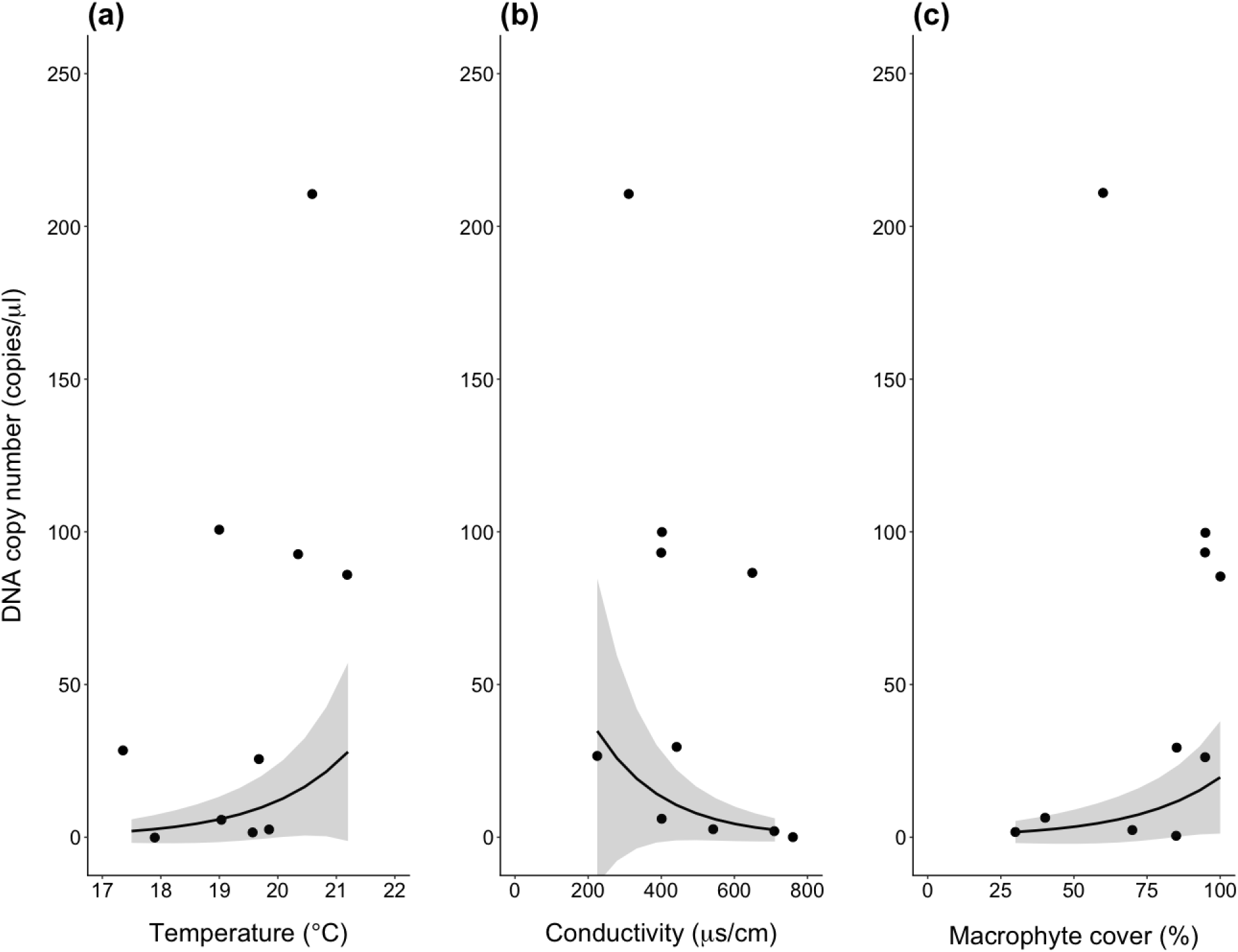
Relationship between fixed effects and response variable (DNA copy number) in ponds, as predicted by the Poisson GLMM. The 95% CIs, as calculated using the model predictions and standard error for these predictions, are given for each relationship. The observed data (points) are also displayed against the predicted relationships (line). DNA copy number increased with water temperature **(a)**, but decreased as conductivity **(b)** and percentage of macrophyte cover **(c)** increased.

### 3.6 PCR versus qPCR

Crucian carp eDNA was amplified by PCR in all samples that amplified using qPCR (Table 2). PCR also provided semi-quantitative estimates of eDNA concentration when PCR products for eDNA samples were run on gels alongside qPCR standards (Figure 6).

**Table 2.**
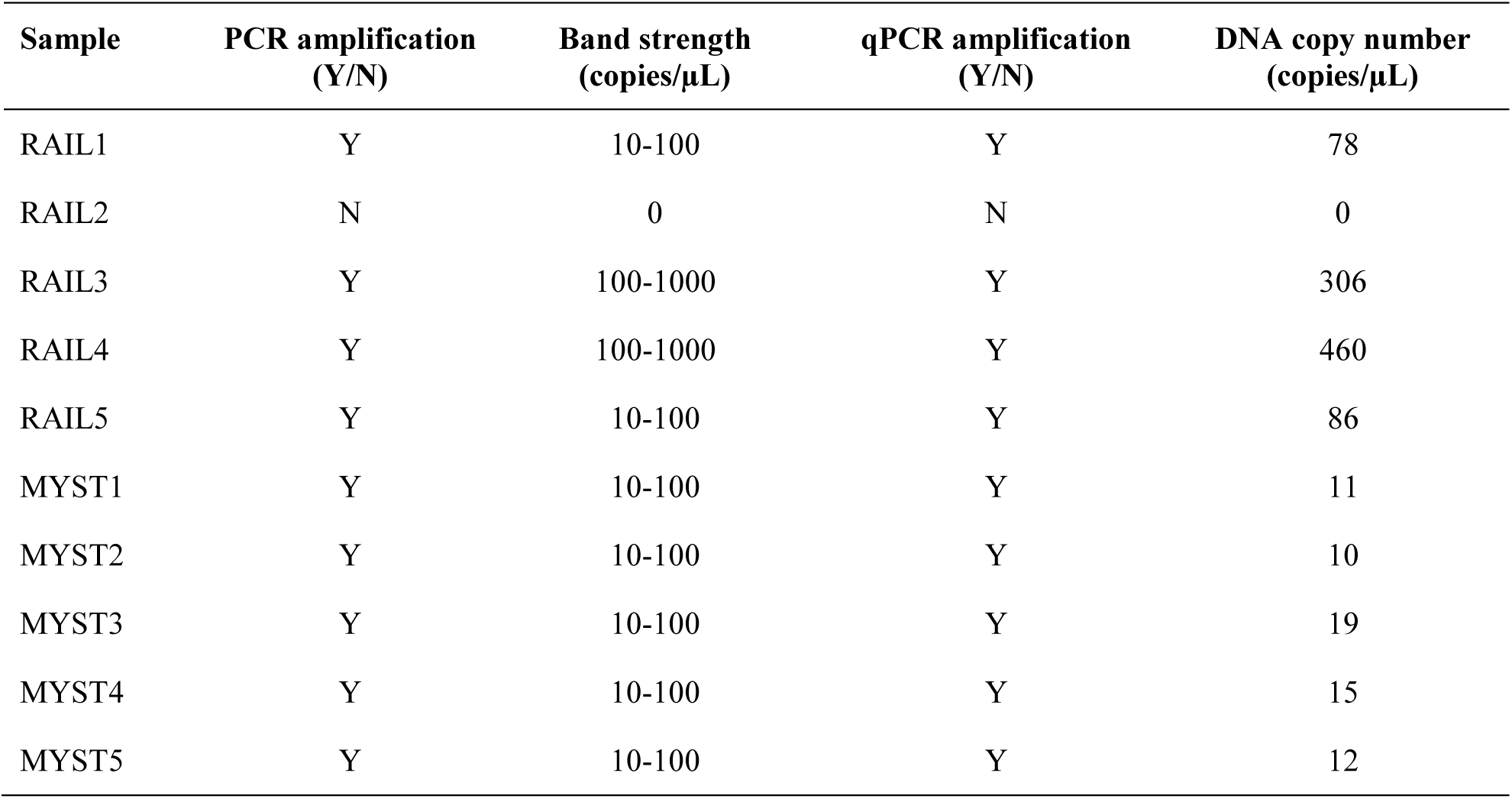
Summary of eDNA amplification by PCR and qPCR for all samples from two ponds.

| Sample | PCR amplification<br>(Y/N) | Band strength<br>(copies/μL) | qPCR amplification<br>(Y/N) | DNA copy number<br>(copies/μL) |
| --- | --- | --- | --- | --- |
| RAIL1 | Y | 10-100 | Y | 78 |
| RAIL2 | N | 0 | N | 0 |
| RAIL3 | Y | 100-1000 | Y | 306 |
| RAIL4 | Y | 100-1000 | Y | 460 |
| RAIL5 | Y | 10-100 | Y | 86 |
| MYST1 | Y | 10-100 | Y | 11 |
| MYST2 | Y | 10-100 | Y | 10 |
| MYST3 | Y | 10-100 | Y | 19 |
| MYST4 | Y | 10-100 | Y | 15 |
| MYST5 | Y | 10-100 | Y | 12 |

**Figure 6.**
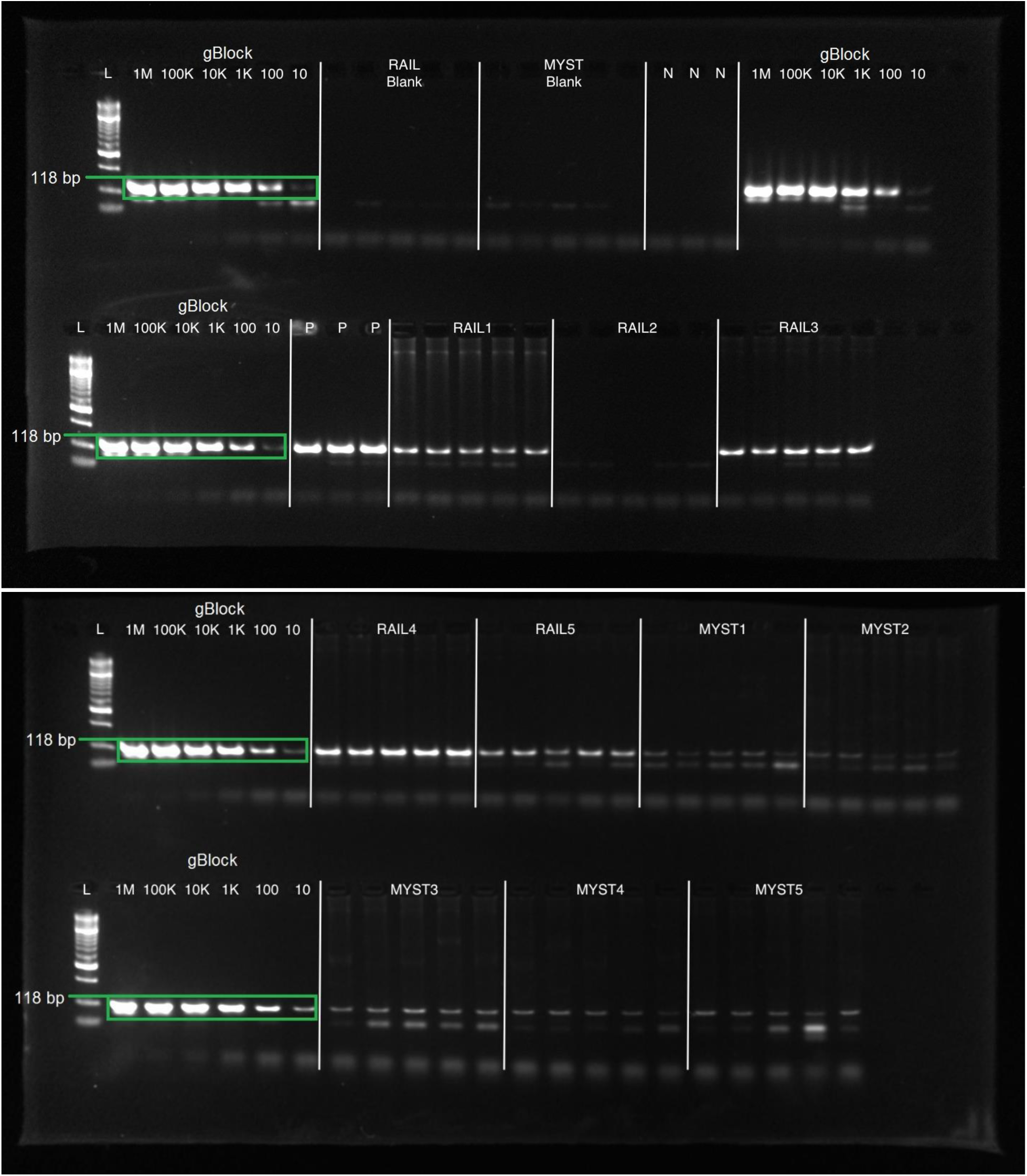
PCR products of gBlocks^®^ standards and five eDNA samples from two ponds. Products were run on 2% agarose gels with Hyperladder™ 50bp (Bioline®, London, UK) molecular weight marker (L). Five replicates were performed for each standard curve point and each eDNA sample. Sample ID is given for each set of replicates, confined by white lines. Exemplary bands of expected size (118 bp) are highlighted in green.

## 4. Discussion

We developed a novel species-specific qPCR assay to enable large-scale distribution monitoring of the threatened crucian carp using eDNA. Crucian carp were detected at almost all sites with confirmed presence and no false positives were generated. Our eDNA approach may have limited suitability for abundance estimation as DNA copy number did not correlate with crucian carp density estimated from netting. However, several biotic and abiotic factors that influence eDNA detection and quantification were identified. Finally, PCR provided semi-quantitative estimates of eDNA concentration and may be a viable alternative to qPCR where funding or laboratory facilities are limited. We discuss areas for improvement in our workflow and provide recommendations for future study.

### 4.1 Using eDNA for crucian carp conservation

eDNA analysis is often compared to conventional monitoring tools to assess performance, reliability, reproducibility, and prospective applications in conservation programmes. We found strong agreement between eDNA and fyke netting for crucian carp detection, where eDNA detected crucian carp in 90% of ponds with presence confirmed by netting. This high detection and low false negative rate supports applicability of eDNA to crucian carp presence-absence monitoring, particularly at large spatial scales where fyke netting is too costly and time-consuming. Abundance estimation is less straightforward as DNA copy number did not directly correspond to crucian carp density. This inconsistency is more likely to stem from eDNA than fyke netting due to effects exerted by external factors (section 4.2) and sample processing (section 4.3) on eDNA quality. However, fyke netting also has detection biases that may influence performance comparisons with eDNA. Fyke net surveys are restricted spatially and temporally (to pre- and post-spawning, as well as spring and autumn when temperatures are low to reduce fish stress in nets), and may fail to capture species that do not have homogenous distribution in their environment (Turner et al., 2012). Netting can be biased towards a particular sex and size class, and catchability dependent on time of year (Ruane, Davenport & Igoe, 2012) and even time of day (Hardie, Barmuta & White, 2006). Therefore, effectiveness of standard methods must also be evaluated and eDNA compared to multiple tools before deemed capable or incapable of estimating abundance.

### 4.2 Factors influencing eDNA detection and quantification

Effects of biotic and abiotic factors on eDNA may vary across target species and ecosystems (Barnes et al., 2014). We found that macrophyte cover negatively influenced eDNA occurrence in ponds, but positively influenced DNA copy number. Crucian carp prefer ponds with stands of aquatic vegetation; however, target DNA may experience interference from plant DNA during qPCR or qPCR reactions may be inhibited by substances in plants, impairing eDNA detection (Jane et al., 2015; Stoeckle et al., 2017). Whilst aquatic vegetation may impede eDNA detection, it may facilitate eDNA preservation and accumulation through reduced UV exposure or induced anoxia (Barnes et al., 2014; Pilliod et al., 2014; Weltz et al., 2017).

eDNA detection probability in qPCR replicates increased at higher crucian carp densities, but decreased as conductivity increased. DNA copy number and conductivity were also negatively correlated. Density is frequently reported to improve detection probability of aquatic species due to more eDNA deposition in the environment (Schmelzle & Kinziger, 2016; Buxton et al., 2017b; Stoeckle et al., 2017). Conductivity has been suggested to influence eDNA detection and quantification, but studies that directly measured this variable found no discernable effect (Takahara et al., 2012; Keskin, 2014; Goldberg et al., 2018). Conductivity (also measured as TDS) relates to sediment type which influences eDNA detection probability, the rate at which sediment binds eDNA, and release of inhibitory substances (Buxton et al., 2017a; Stoeckle et al., 2017). Notably, the only false negative pond in our study was also the most conductive (760 μs/cm). Therefore, conductivity may lead to incorrect inferences about species presence and impact conservation management decisions.

Our results indicate that samples may have been affected by inhibitory substances despite tests performed to identify inhibition. We spiked qPCR reactions with a known amount of synthetic target DNA. However, an artificial Internal Positive Control gene assay may identify inhibition more effectively (Goldberg et al., 2016). Dilution of eDNA samples (and inhibitory substances present) can release inhibition, but also reduce detection probability (Piggott, 2016) and induce false negatives (Buxton et al., 2017a). We used TaqMan^®^ Environmental Master Mix 2.0 (Life Technologies, CA, USA) in qPCR reactions to counter inhibition (Jane et al., 2015), but it may be advisable to use DNA extraction kits that perform inhibitor removal (Sellers et al., 2018) or include Bovine-serum albumin (BSA) in qPCR reactions (Jane et al., 2015). Alternatively, ddPCR may handle inhibitors better than qPCR and provide more accurate abundance or biomass estimates (Nathan et al., 2014).

In addition to effects of macrophyte cover and conductivity, water temperature positively influenced DNA copy number. Although warmer temperature coincided with breeding activity and heightened DNA release in other fish and amphibian species (Buxton et al., 2017b; Bylemans et al., 2017), water sample collection in late August was outwith the reported spawning period for crucian carp (Aho & Holopainen, 2000). The association observed here may instead reflect increased DNA shedding rates caused by higher metabolic activity in response to warm temperature, as reported for other fish species (Takahara et al., 2012; Robson et al., 2016; Lacoursière-Roussel et al., 2016).

Crucially, environmental data were not collected in 2016 for every pond in our study. Our results indicate direction of effects of biotic and abiotic factors on eDNA detection and quantification, but contemporary data (particularly temperature) are needed for comprehensive interpretation of these relationships. However, it is clear that eDNA practitioners must account for these effects as well as sample inhibition. The uncertainty around the estimated effects of covariates at each level of our hierarchical occupancy model and GLMM also imply that greater sample volume, sample number, and/or qPCR replication are required to improve the ability and precision of our assay to detect crucian carp eDNA and reduce the potential for false negatives (Schultz & Lance, 2015; Goldberg et al., 2018).

### 4.3 Optimisation of eDNA workflow

Some non-target DNA extracts used to validate assay specificity were contaminated with crucian carp DNA. Field cross-contamination can occur if reference tissue material is collected from multiple species without sterilising equipment, or eDNA is present on the material collected (Rodgers, 2017). Collection and storage of reference tissue material is an important consideration for eDNA practitioners, particularly those using highly sensitive assays (LOD = 1 copy/μL) (Wilcox et al., 2013, 2016). Dedicated, sterilised equipment should be used when collecting new reference material from different species. From existing reference collections, only non-target samples that were collected on separate and chronologically distinct occasions from target samples should be used (Rodgers, 2017).

Cross-contamination can also arise during water sampling, filtration, DNA extraction and qPCR preparation. Low-level contamination was found in one full process blank but detections from this pond were not omitted as it contained crucian carp and contamination was not observed downstream. All equipment in our study was sterilised by immersion in 10% chlorine-based commercial bleach solution for 10 mins, followed by 5% MicroSol detergent (Anachem, UK), and rinsed with purified water. However, sterilisation with 50% chlorine-based commercial bleach solution (Goldberg et al., 2016) or single-use, sterile materials (Wilcox et al., 2016) may further minimise contamination risk.

Many of our eDNA samples were low concentration (<100 copies/µL) which can cause inconsistent qPCR amplification (Goldberg et al., 2016), thus we discuss approaches to maximise eDNA concentration and improve detection probability. The probability of eDNA detection depends heavily on the number of samples and volume of water collected, time of sampling, and sample concentration (Schultz & Lance, 2015; Goldberg et al., 2018). We sampled 5 × 2 L water samples from each pond in autumn 2016, but timing and/or sampling effort may have been inappropriate. A seasonal effect on common carp eDNA detection was observed, where spring sampling generated higher eDNA concentration and detection rate due to greater common carp activity (Turner et al., 2014) and density (Hinlo et al., 2017a). As water sampling did not coincide with fyke netting (spring 2016) in our study, eDNA concentration may not reflect CPUE estimates. Water samples in spring may contain more crucian carp eDNA due to higher activity of individuals, or autumn fyke netting may produce lower CPUE estimates. Parallel seasonal sampling, where water sampling is performed in conjunction with fyke netting throughout the year, may better align eDNA concentration with CPUE estimates and enable eDNA-based abundance estimates for crucian carp.

Representative sampling is crucial in eDNA surveys. Individuals of a species can be unevenly distributed in the environment, which impacts eDNA detection, distribution, and concentration (Takahara et al., 2012; Eichmiller, Bajer & Sorensen, 2014; Schmelzle & Kinziger, 2016; Goldberg et al., 2018). In lentic ecosystems, eDNA has a patchy horizontal and sometimes vertical distribution, resulting in fine spatial variation (Eichmiller et al., 2014). Studies on common carp revealed eDNA was more concentrated near the shoreline of lentic water bodies (Takahara et al., 2012; Eichmiller et al., 2014), due to aggregations of individuals (Eichmiller et al., 2014). We collected surface water (5 × 2 L) from the shoreline and sampled at equidistant points around the pond perimeter where possible; however, more samples and greater water volumes may be required to improve detection probability (Schultz & Lance, 2015; Goldberg et al., 2018). Fine spatial sampling and occupancy modelling are needed to determine the sample number required to achieve high detection probability and eliminate false negatives (Goldberg et al., 2018). However, the number of samples required will inevitably vary across habitats due to inherently variable physical properties (Schmelzle & Kinziger, 2016).

Method of eDNA capture can dictate success of this monitoring tool. Studies of eDNA in ponds (Ficetola et al., 2008; Biggs et al., 2015) have used an ethanol precipitation approach, but this is restricted to small volumes. Filtration allows more water to be processed and minimises capture of non-target DNA, with macro-organism eDNA effectively captured by pore sizes of 1 - 10 μm (Turner et al., 2014). We used a small pore size of 0.45 μm to capture most eDNA particle sizes, although filter clogging prevented the entire sample being processed and may have affected eDNA concentration downstream. Pre-filtering can reduce clogging, but is labour-intensive and increases cost (Takahara et al., 2012). Larger pore sizes have been used in temperate and tropical lentic environments (Takahara et al., 2012; Robson et al., 2016; Goldberg et al., 2018), though independent investigation is needed to determine which pore size maximises target DNA concentration.

Comparisons of eDNA yield across filter types and DNA extraction protocols have shown that cellulose nitrate filters stored at −20 °C (this study) often provide best eDNA yield (Piggott, 2016; Spens et al., 2016; Hinlo et al., 2017b). However, different filter types may be optimal for different species, which has consequences for detectability (Spens et al., 2016) and relationships between eDNA concentration and abundance/biomass (Lacoursière-Roussel et al., 2016). Extraction method used, regardless of filter type, will ultimately influence DNA quality and yield. We used the PowerWater^®^ DNA Isolation Kit (MO BIO Laboratories, CA, USA), but the DNeasy Blood and Tissue kit (Qiagen^®^, Hilden, Germany) has demonstrated greater yield (Hinlo et al., 2017b). We also combined filters from the same sample for DNA extraction at the bead milling stage, but independent lysis may recover more DNA (Hinlo et al., 2017b). A comparison of DNA extraction protocols is necessary to assess which maximises crucian carp eDNA concentration. A new modular extraction method shows promise for eDNA but has yet to be evaluated for targeted qPCR (Sellars et al., 2018).

Finally, detection sensitivity can be enhanced by increasing the number of qPCR technical replicates (Schultz & Lance, 2015; Piggott, 2016). We performed five technical replicates for each of our samples, but other studies have used as many as twelve and only one may amplify (Biggs et al., 2015). More replication may have enabled amplification from CHIP samples, but qPCR cost would inevitably increase. Further replication may also be unnecessary if steps are taken to improve initial concentration of samples instead (Schultz & Lance, 2015).

### 4.4 PCR or qPCR?

Our study is not the first to compare eDNA detection using different means of DNA amplification (Nathan et al., 2014; Farrington et al., 2015; Piggott, 2016; De Ventura et al., 2017). Like Nathan et al. (2014), we found PCR had comparable sensitivity to qPCR and band strength of PCR products may indicate eDNA concentration; however, we also translated band strength to approximate DNA copy number. PCR may require more replication to achieve set detection probabilities (Piggott, 2016), but lower sensitivity could make this approach more robust to false positives from cross-contamination than qPCR (De Ventura et al., 2017). Large-scale comparisons of PCR and qPCR across study systems and species are needed to truly assess performance of each approach. Nonetheless, our findings support PCR as a cost-efficient, semi-quantitative alternative to qPCR for conservation programmes wishing to utilise eDNA (Nathan et al., 2014; De Ventura et al., 2017).

### 4.5 Concluding remarks

A primary objective of the Norfolk crucian carp BAP was to obtain a basic understanding of species distribution and population status across Norfolk (Copp & Sayer, 2010). eDNA surveillance for crucian carp will provide a useful, cost-effective alternative to established survey methods where the aim is determining presence-absence. Our assay may detect hybrids where crucian carp were the maternal parent due to use of a mitochondrial marker; however, these detections are also beneficial to the crucian carp conservation effort by identifying ponds where true crucian carp may still exist and contamination with goldfish, common carp and their hybrids has occurred. Alternatively, our assay could be used as an early warning tool in countries where the crucian carp is considered invasive. The areas we have highlighted require further investigation before eDNA can be used routinely. Nevertheless, eDNA survey could enable large-scale distribution monitoring for crucian carp through rapid screening of existing and new ponds. Fyke netting could then be used to investigate age, sex and size structure of populations, and remove hybrids.

## Acknowledgements

We would like to thank landowners and land managers for granting permission to sample the ponds included in this study, and Ian Patmore, Dave Emson, Helen Greaves and Glenn Wiseman for assistance with fyke netting. We are grateful to Marco Benucci for assistance with water sampling and filtration, Graham Sellers for advice on primer design and validation, and Peter Shum for support with qPCR troubleshooting and feedback on the manuscript.

